# A unique case in which *Kimoto*-style fermentation was completed with *Leuconostoc* as the dominant genus without transitioning to *Lactobacillus*

**DOI:** 10.1101/2022.09.05.505584

**Authors:** Kohei Ito, Ryo Niwa, Yuta Yamagishi, Ken Kobayashi, Yuji Tsuchida, Genki Hoshino, Tomoyuki Nakagawa, Takashi Watanabe

## Abstract

The *Kimoto*-style fermentation starter is a traditional preparation method of *sake* brewing. In this process, specific microbial transition patterns have been observed within nitrate-reducing bacteria and lactic acid bacteria during the production process of the fermentation starter. We have characterized phylogenetic compositions and diversity of the bacterial community in a *sake* brewery performing the *Kimoto*-style fermentation. Comparing the time-series changes with other *sake* breweries previously reported, we found a novel type of *Kimoto*-style fermentation in which the microbial transition differed significantly from other breweries during the fermentation step. Specifically, the lactic acid bacteria, *Leuconostoc* spp. was a predominant species in the late stage in the preparation process of fermentation starter, on the other hand, *Lactobacillus* spp., which plays a pivotal role in other breweries, was not detected in this analysis. The discovery of this new variation of microbiome transition in *Kimoto*-style fermentation has further deepened our understanding of the diversity of *sake* brewing.

## INTRODUCTION

*Sake* is a traditional alcoholic beverage made from rice in Japan. In a preliminary stage of a main brewing process, a fermentation starter with purely cultured yeast is produced to prevent microbial contamination and poor fermentation and to promote smooth alcoholic fermentation. The production process is as follows; First, *Aspergillus oryzae*, which secretes amylases, is propagated on steamed rice to make *koji*. Then, *koji*, steamed rice, and water are mixed in an open-top tank. Fermentation of this mixture produces a fermentation starter. The fermentation starter is further mixed with *koji*, steamed rice, and water, and after a 3-5 week fermentation process, the fermentation mash or starter is produced. The fermentation starter is separated into *sake* and spent rice by a filter press to complete the *sake*.

The fermentation starter is divided into three styles, *Sokujo, Kimoto* and *Yamahai. Sokujo*-style fermentation starter is a modern method to make the starter culture with the addition of food-grade lactic acid. *Kimoto*-style fermentation starter is the traditional preparation method of the starter culture and is manufactured under highly acidic conditions by inducing the growth of nitrate-reducing bacteria and lactic acid bacteria properly. *Yamahai*-style is similar to *Kimoto*-style but made without grinding rice. Lactic acid inhibits contaminations of unintended yeasts and bacteria from external environments into the fermentation starter.

The microbiome compositions during *Kimoto*-style fermentation starter production show standard transitions as follows. Nitrate-reducing bacteria, which were reported to come from water (1), initially grows and produces nitrite, thereby inhibiting the growth of microorganisms that are less tolerant to nitrite. At the same time, lactic acid bacteria, especially *Leuconostoc* spp., which grow at low temperatures and have fewer nutrient requirements, increase, and then lactic acid bacteria such as *Latilactobacillus sakei*, which require strict nutrients, occupy the microbiome compositions as a predominant species. These steps are known as a common microbial transition in *sake* brewing (2–4).

However, it was reported that some fermentation starters brewed by *Kimoto*-style in several *sake* breweries show distinctive microbial transitions and chemical changes and that this is one of the reasons for producing unique *sake* flavors among breweries, even if they use the same production process (5–8). A possible reason to explain this fact is differences in *Kuratsuki* microorganisms (microorganisms living in *sake* breweries) and the introduction of diverse microorganisms from outside of tanks during the *sake* brewing process (9,10).

The *Tsuchida Sake* Brewery (Gunma, Japan) is one of a few breweries that produce the *Kimoto*-style fermentation starter without adding yeasts and fully relies on *Kuratsuki* microorganisms to produce the fermentation starter. In addition, compared to the *Kimoto*-style fermentation in other *sake* breweries, this *sake* brewery is characterized by not using any food additives such as brewers’ alcohol, enzyme reagents, or activated charcoal. Furthermore, this brewery does not use *sake*-brewing rice but table rice for *sake* brewing. For the above reasons, the microbial community, and its transition in the fermentation starter of the brewery were considered to be divergent from previous studies.

In this study, we focused on the fermentation mechanism of the *Kimoto*-style fermentation starter from the *Tsuchida Sake* Brewery and analyzed its microbial community during the fermentation process in detail using its 16S ribosomal RNA amplicon sequencing. The *Kimoto*-style fermentation starter from the *Tsuchida Sake* Brewery possesses a distinctive microbial community compared with other *Kimoto*-style fermentation starters; specifically, the dominant genus of lactic acid bacteria was *Leuconostoc*, and the switch of the predominant lactic acid bacteria, from *Leuconostoc* to *Lactobacillus*, was not observed in the fermentation process of the *Kimoto*-style fermentation starter. Here we report a new profile of microbial transition in the *Kimoto*-style fermentation starter.

## MATERIALS AND METHODS

### Sample collection

Samples were collected for each day 1, 3, 5, 7, 9, 13, 17, 22, 28, and 33 after starting the fermentation of the fermentation starter. All samples were collected in between October and November 2021. The ingredients of the fermentation starter were water, polished rice, and *koji*. We did not use nitric acid or other additives for this batch. The polishing ratio of the rice was 85%. It is assumed that the microbiome is not homogeneous because the fermentation starter is produced in a large tank. Therefore, all samples were collected in duplicate to avoid sampling bias. Fermentation starter samples used in this study were provided by *Tsuchida Sake* Brewing Company (Gunma, Japan). All samples were immediately frozen and stored until DNA extraction.

### Measurement of basic chemical components

Temperatures and chemical compositions of the fermentation starter were measured. Alcohol (ALC) and Baumé degree (Be) were measured using a DA-155 vibrating density meter for alcoholic beverages (Kyoto Electronics Manufacturing Co., Ltd., Kyoto, Japan), while Acidity (TA) and Amino Acid Content (AA) were obtained using a CHA-700 multi-sample changer (Kyoto Electronics Manufacturing Co., Ltd.). The nitrite concentration was measured using a Merck Millipore MQuant nitrite test (Merck KGaA, Darmstadt, Germany).

### Total DNA extraction and high throughput sequencing

Samples were subjected to 750 μl of lysis buffer supplied with the GenFind V2 DNA extraction kit (Beckman Coulter, Indianapolis, U.S.). The suspension was vortexed for 10 min, heat-treated at 100 °C for 10 min, and centrifuged for 5 min at 15,000 rpm. The supernatant was mixed with E.Z. beads (AMR, Tokyo, Japan). DNA fragmentation was performed on an MM-400 (Retsch, Haan, Germany) at a maximum speed of 3 min. The rest of the DNA purification process was performed using a GenFind v2 DNA extraction kit (Beckman Coulter, Indianapolis, U.S.) following the manufacturer’s protocol. Finally, DNA was eluted with 80 μl of sterile water. 341F (5’-TCGTCGGCAGCGTCAGATGTGTATAAGAGAGACACCTACGGGNGGCWGCAG-3’) and 806R (5’-GTCTCGTGGGCTCGGGAGATGTGTATAAGAGACAGGACTACHVGGGTATCTAATC C-3’) primers were utilized to amplify the V3-V4 region of the 16S rRNA gene by PCR (11,12). Thermal cycling conditions were 95 °C for 3 min; 32 cycles of 95 °C for 30 s, 55 °C for 30 s, and 72 °C for 30 s; and a final extension at 72 °C for 5 min. A second PCR was performed to add sequencing adapters and dual-index barcodes to distinguish amplicons from each sample using the same reaction conditions with eight cycles. Preparation of libraries and sequencing were performed by paired-end sequencing of 300 bp on the Illumina MiSeq platform (Illumina, Inc., San Diego USA) at GenomeRead Inc. in Kagawa, Japan.

### Microbiome analysis

The QIIME2 (version 2021.02) (13) platform was used for microbiome analysis. FASTQ files were imported into the QIIME2 platform. Sequences were processed using qiime dada2 denoise-paired command for quality control and classification into amplicon sequencing variants (ASVs). Taxonomic analysis was run on SILVA database SSU 138 by qiime feature-classifier classify-sklearn (14–16). Sequence reads that were not classified as any species after phylogenetic classification (Unassigned) and reads classified as chloroplast and mitochondria were excluded from further analyses. Since the genus *Latilactobacillus* is a taxon which relatively recently derived from the genus *Lactobacillus* (17), the SILVA database SSU 138 used in this study does not reflect the reclassification of the genus *Latilactobacillus*, so it was assigned as *Lactobacillus* in this study.

### Processing for comparative microbiome analysis

Sequence data from BioProject PRJDB12939 was used for comparative analysis (8). Since 341F (5’-CCTACGGGGNGGCWGCAG-3’) and 805R (5’-GACTACHVGGGGTATCTAATCC-3’) were used in the study, removal of one base from the 5’ end of the reverse read was performed to match the read lengths of the samples obtained in our study. The fastp v.0.20.1. (18) was deployed for this processing. The method described above was repeated to perform taxonomic analysis. The depth of sequence reads (Features) differed among samples, and to normalize them, we subsampled from each sample to 5,000 reads each. Samples with fewer than 5,000 reads (S22, 23, 24, 25, 26, 28) were excluded from the diversity analysis (Table S1). Sub-sampling is an approach for inferring microbiome differences between samples and has been reported to be a suitable analytical method when analyzing new data sets (19). To evaluate the effect of sequence read counts on microbiome diversity assessment, we examined changes in the value of the Shannon index over a range of reading counts from 0-5,000 by rarefaction curves. The Rarefaction curve of the Shannon index leveled off when the number of leads was just under 500 (Table S2). Therefore, this investigation suggests that no significant changes in microbial diversity are due to the subsampling of 5,000 reads from each sample.

### Statistical Analysis

Kruskal-Wallis tests were used to compare the alpha diversity (Shannon diversity index) among samples. To compare differences in Beta diversity (Weighted UniFrac distance) between samples, for all PERMANOVA analyses, 5,000 trials were performed to assess the statistical significance. Q-values < 0.05 after multiple testing corrections were considered statistically significant. All multiple testing corrections were performed by computing FDRs using the Benjamini–Hochberg method, and Q-values (adjusted P-values) < 0.05 were considered statistically significant. Data visualization was performed by R version 4.2.1, the ggplot2 package version 3.4.0 and ggprism version 1.0.4.

## RESULTS

### Chemicals and Temperature Change in fermentation starters

The chemical data and the average temperature of the *sake* starter are shown in Fig. 1. The amount of nitrite started to rise on day 7 and reached its peak on day 11. On day 14 it was undetectable; Titratable acidity (TA), a value representing total acidity, increased on day 15, followed by an increase in ethanol concentration on day 27. In addition, a rapid decrease in glucose was observed along with an increase in ethanol concentration on day 27. These changes in chemical data and the average temperature are generally observed in *Kimoto*-style fermentation, so we suggest that the fermentation proceeded well in the *Kimoto*-style fermentation starter of the *Tsuchida Sake* Brewery.

**FIG. 1.**
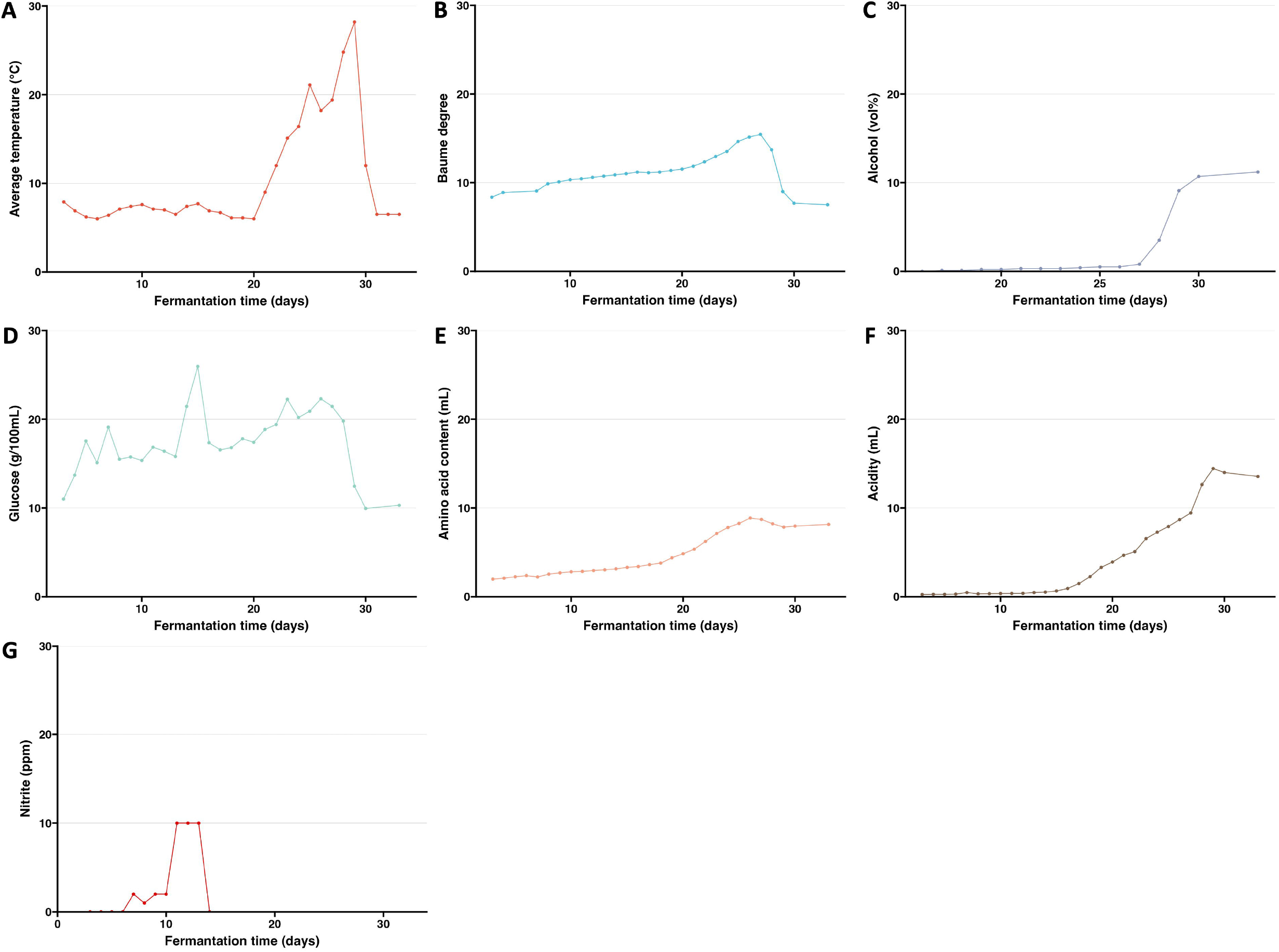
Time-series data of chemical concentrations, degree, and average temperature. Average temperature (A), Baume degree (B), Alcohol (C), Glucose (D), Amino acid content (E), Acidity (F), Nitrite (G).

### Phylogenetic composition of the microbiome found in the fermentation starter

To track the microbial transition in the fermentation starter during the fermentation, we performed 16S rRNA amplicon sequencing. Results demonstrated large changes in relative abundances of the bacterial genera (Genus) during the early to late stages (Fig. 2). Taxonomic compositions of day 7 were markedly different between duplicates.

**FIG. 2.**
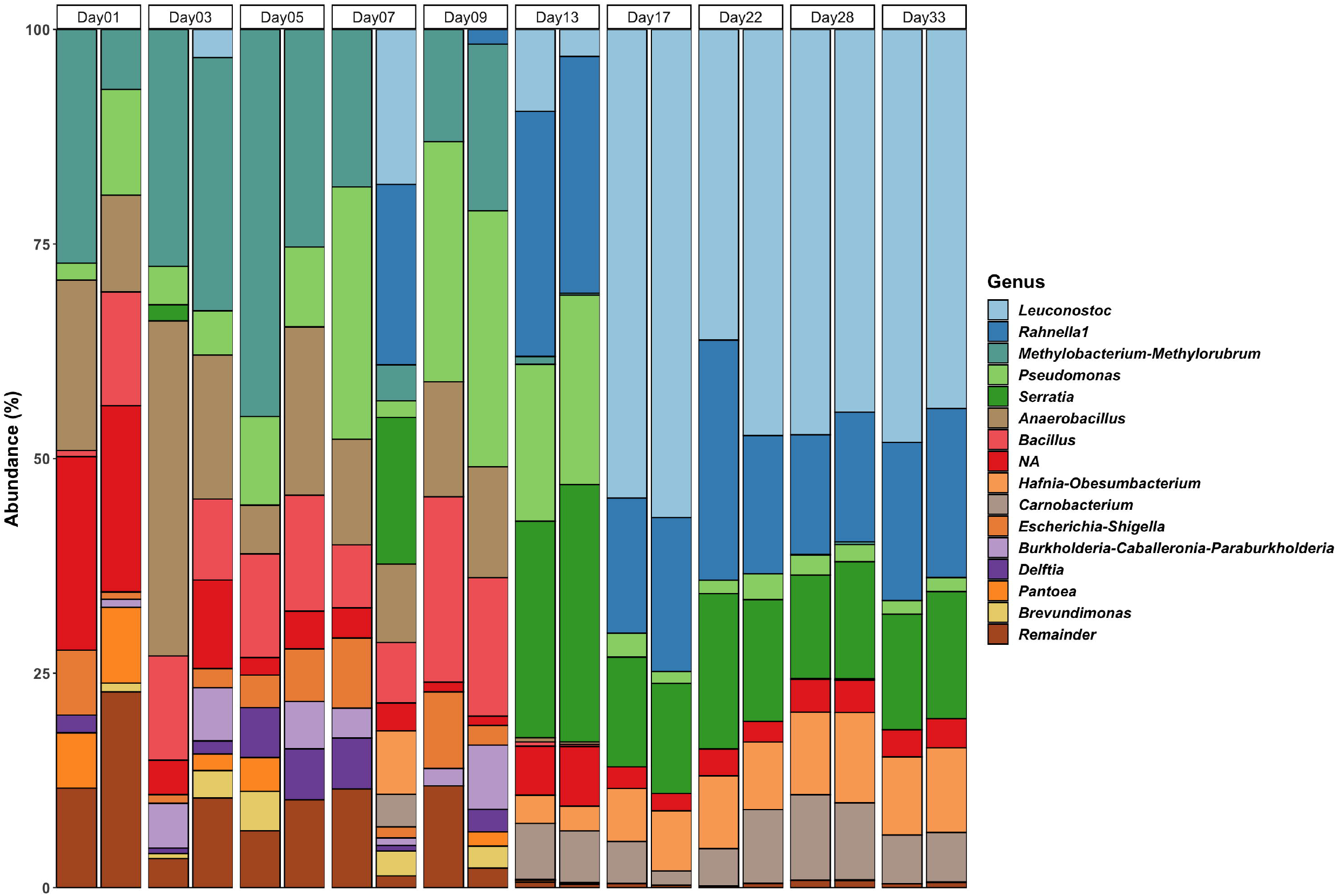
Taxonomic composition of microbiomes during *Kimoto*-style fermentation starter in *sake* brewing. The top 15 genera are listed, and the rest are noted as “Remainder”.

The genera with average relative abundances higher than 5% in the early stage (day 1 to day 7) of the fermentation starters were *Methylobacterium-Methylorubrum* (23.0%), *Anaerobacillus* (16.7%), *Bacillus* (9.4%), and *Pseudomonas* (9.3%) (Table S3). The relative abundances of these bacterial genera decreased significantly within days 7-9. *Anaerobacterium* spp. is salt-tolerant and halophilic and has been found in lakes and soils in arid regions (20,21). It is reported that *Bacillus* spp. may live in *sake* breweries as *Kuratsuki* bacteria (22).

The bacterial genera with average relative abundances higher than 5% in the late stage (days 17-33) were *Rahnella1* (18.1%), *Serratia* (13.9%), *Hafnia-Obesumbacterium* (8.6%), and *Carnobacterium* (6.2%) (Table S3). *Leuconostoc, Rahnella, Serratia, Hafnia-Obesumbacterium*, and *Carnobacterium* were the bacterial genera that accounted for a large proportion of the bacteria found in the late stage (days 17-33). *Leuconostoc* spp. has been detected in several fermented foods (23,24).

### Comparison with other breweries

To characterize the microbial transition in this brewery, we compared our data with a previous study that investigated the microbial transition in four *sake* breweries (8). Time-series changes in the microbial diversity (Alpha diversity; Shannon index) for each sample are shown (Table 1). *Tsuchida Sake* brewery was confirmed to have a higher microbial diversity (Shannon index) of the fermentation starters (Q-value < 0.05) compared to the other breweries (Table S4).

**TABLE 1.** Time-series changes in Shannon index for each sample.

| days | sample_type | brewery | Shannon index |
| --- | --- | --- | --- |
| 5 | shubo | Tsuchida | 4.083626084 |
| 7 | shubo | Tsuchida | 4.483739465 |
| 9 | shubo | Tsuchida | 4.00914674 |
| 9 | shubo | Tsuchida | 3.260447879 |
| 13 | shubo | Tsuchida | 3.955204079 |
| 13 | shubo | Tsuchida | 3.953384193 |
| 17 | shubo | Tsuchida | 2.683731987 |
| 17 | shubo | Tsuchida | 2.54526448 |
| 22 | shubo | Tsuchida | 3.246816955 |
| 22 | shubo | Tsuchida | 2.970864866 |
| 28 | shubo | Tsuchida | 2.966174624 |
| 28 | shubo | Tsuchida | 3.072856664 |
| 33 | shubo | Tsuchida | 2.925714709 |
| 33 | shubo | Tsuchida | 3.056336844 |
| 3 | shubo | A | 0.075588383 |
| 6 | shubo | A | 0.47124225 |
| 9 | shubo | A | 1.748384977 |
| 10 | shubo | A | 1.517717158 |
| 11 | shubo | A | 1.252412288 |
| 12 | shubo | A | 1.496213641 |
| 13 | shubo | A | 1.682703746 |
| 15 | shubo | A | 1.732575907 |
| 16 | shubo | A | 1.486121819 |
| 3 | shubo | B | 2.38193983 |
| 6 | shubo | B | 2.933967722 |
| 7 | shubo | B | 3.090082101 |
| 8 | shubo | B | 3.294985987 |
| 9 | shubo | B | 3.142318413 |
| 10 | shubo | B | 2.2285432 |
| 12 | shubo | B | 1.835421019 |
| 13 | shubo | B | 2.113163833 |
| 14 | shubo | B | 2.012528982 |
| 3 | shubo | C | 1.837678435 |
| 4 | shubo | C | 2.042470055 |
| 5 | shubo | C | 1.28555966 |
| 7 | shubo | C | 1.261663511 |
| 8 | shubo | C | 1.221165987 |
| 9 | shubo | C | 1.037990141 |
| 11 | shubo | C | 1.10982519 |
| 13 | shubo | C | 1.085215814 |
| 16 | shubo | C | 1.016089562 |
| 3 | shubo | D | 1.812960613 |
| 6 | shubo | D | 2.714883385 |
| 8 | shubo | D | 3.393236556 |
| 9 | shubo | D | 2.492923589 |
| 10 | shubo | D | 2.097860823 |
| 11 | shubo | D | 2.214411678 |
| 13 | shubo | D | 2.233278552 |
| 3 | shubo | D2 | 0.541406715 |
| 5 | shubo | D2 | 0.55204851 |
| 6 | shubo | D2 | 0.460579254 |
| 7 | shubo | D2 | 0.725185898 |
| 9 | shubo | D2 | 2.65814023 |
| 10 | shubo | D2 | 1.727365975 |
| 11 | shubo | D2 | 1.722444728 |
| 12 | shubo | D2 | 1.675929835 |
| 14 | shubo | D2 | 1.64604447 |
| 3 | shubo | E | 0.806481485 |
| 5 | shubo | E | 1.332881408 |
| 6 | shubo | E | 2.553896711 |
| 7 | shubo | E | 1.644847183 |
| 9 | shubo | E | 1.56711282 |
| 11 | shubo | E | 1.427864233 |
| 12 | shubo | E | 1.328343618 |
| 13 | shubo | E | 1.247446865 |
| 16 | shubo | E | 1.325637366 |

A Principal Coordinate Analysis (PCoA) by Weighted UniFrac distance to visualize the differences (Beta diversity) among the microbiomes of each sample showed that the *Tsuchida Sake* Brewery formed a different cluster (q-value < 0.007) than the other breweries (Fig. 3 and Table S5).

**FIG. 3.**
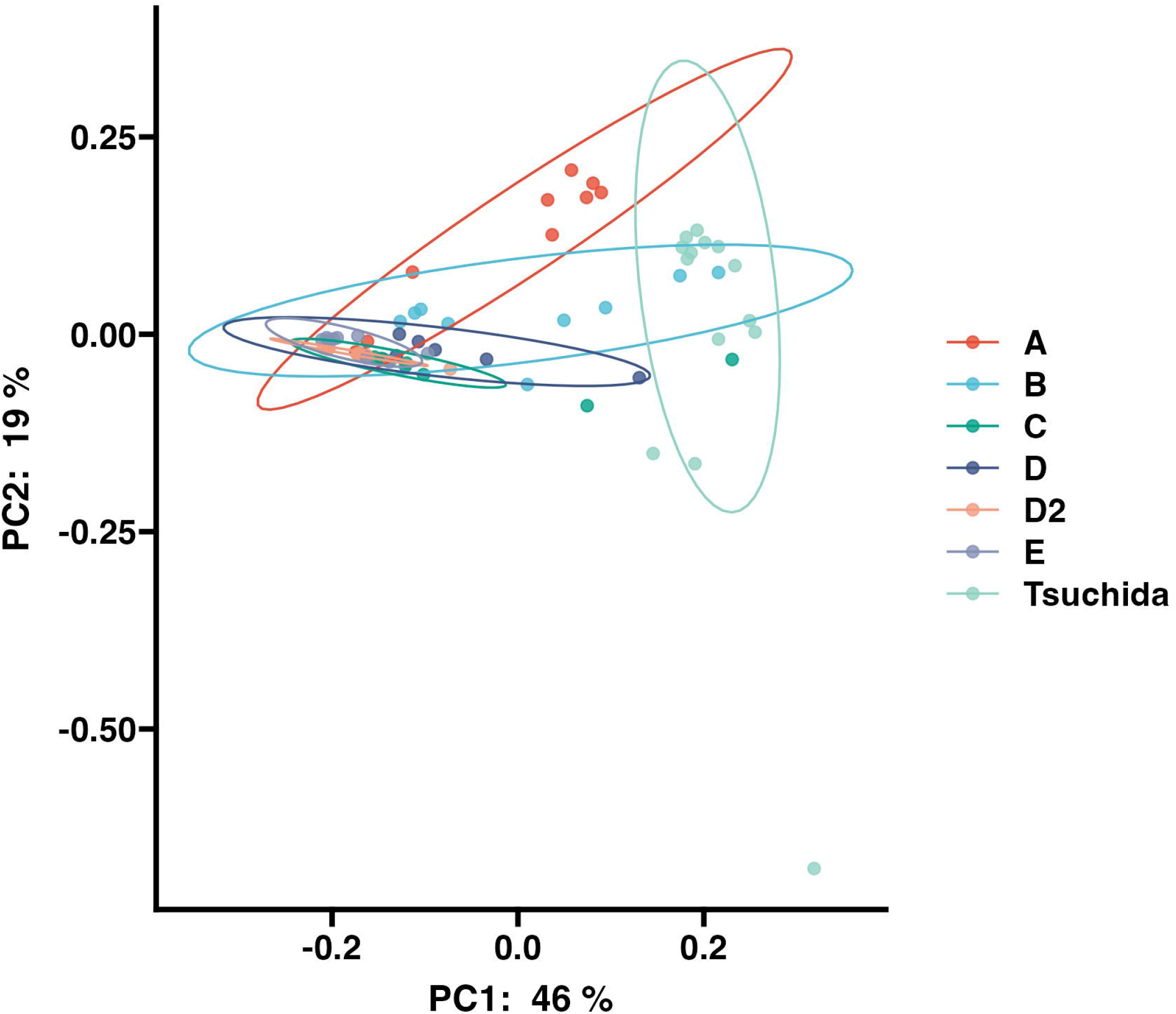
Principal coordinate analysis (PCoA) of 5 *sake* breweries. Samples are compared using the weighted UniFrac distance metrics.

The relative abundances of the genus *Lactobacillus* and *Leuconostoc* in each brewery during the preparation are shown in the line graphs (Fig. 4). The genus *Lactobacillus* became the dominant species in all other breweries, but the relative abundance of the genus *Lactobacillus* did not increase during the entire preparation in the *Tsuchida Sake* Brewery. On the other hand, the genus *Leuconostoc* increased in only three breweries, and in all but the *Tsuchida Sake* Brewery, their abundance eventually decreased and shifted to the genus *Lactobacillus*. In contrast, in the *Tsuchida Sake* Brewery, the genus *Leuconostoc* maintained a constant high relative abundance during the fermentation.

**FIG. 4.**
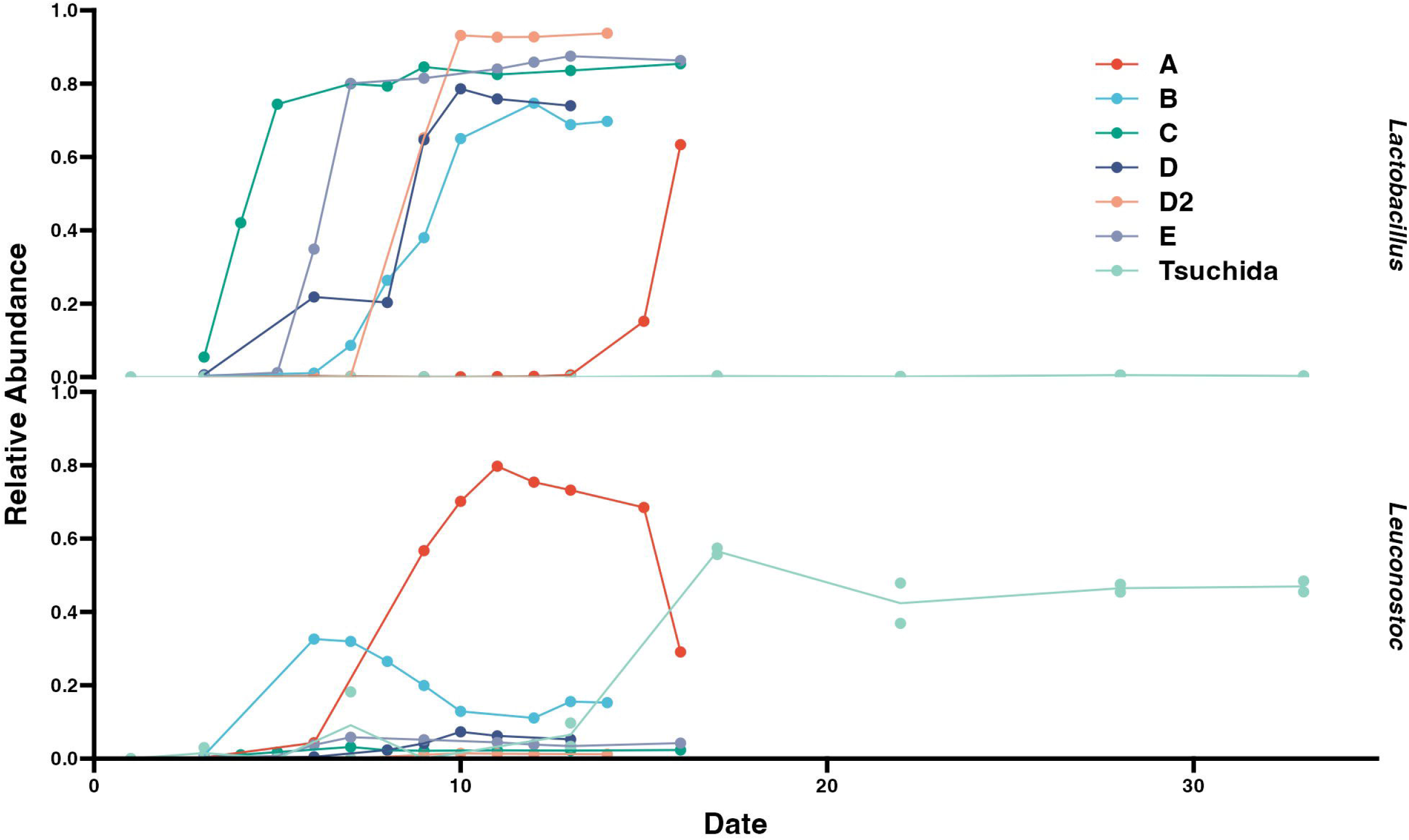
Line plots of relative abundances of *Lactobacillus* and *Leuconostoc*.

## DISCUSSION

### Comparative analysis revealed unique dynamics of lactic acid bacteria in the *Tsuchida Sake* Brewery

A previous study reported three major microbial transition profiles in *sake* brewing from the point of switches occurring in lactic acid bacteria: (1) lactic acid bacteria increase overall, but *Lactobacillus* spp. remain more abundant than *Leuconostoc* spp., (2) *Lactobacillus* spp. are more abundant in the beginning, but *Leuconostoc* spp. become dominant later, and (3) *Leuconostoc* spp. remain more abundant than *Lactobacillus* spp. (8). Surprisingly, in the brewery we sampled, the genus *Leuconostoc* was detected as the dominant species even on day 33, the last day of brewing, and only low abundances of the genus *Lactobacillus* was detected throughout the entire preparation, indicating a microbial transition profile differs from previous reports.

In addition, another study reported that *Kimoto*-style fermentation is characterized by the detection of multiple lactic acid bacteria compared to *Sokujo-moto* (25), so the process of Kimoto-style fermentation itself may contribute to the diversity of taste in different *sake* breweries. *Sokujo-moto* is a modern type of fermentation starter which includes lactic acid to maintain a low pH, thus preventing microbial contamination from the brewery (5). As indicated in this study, specific microbiome transitions were observed that were different from the microbial transitions reported by Takahashi *et al*., suggesting that the microbiome is unique in individual breweries and that these microbial transition factors that characterize them need to be investigated (8).

Some studies reported that only one lactic acid bacteria appears and the transition does not occur (26), and no *Lactobacillus* spp. are found (27), but to the best of my knowledge this is the first study proving this trend of microbiome shift in a manner of uncultured analysis method.

A previous study revealed that *L. sakei* has a more stringent amino acid requirement than *L. mesenteroides, Lactobacillus* sp., and di-tripeptides including asparagine, which is produced by *koji* mold degradation, are growth factors, and that pH and temperature affect the growth of *L. sakei* (8,28,29). In this study, the peak of nitrite concentration was relatively late at day 11, and the increase in acidity (TA; Titratable acidity) was delayed accordingly, suggesting a longer survival period of adventitious bacteria that were initially introduced (Fig. 2). Therefore, there is a possibility that the growth of *L. sakei* was inhibited by specific changes in nutrients and temperature.

In addition, D-amino acids are attracting attention as new taste components of *sake*, D-alanine is reported to be produced by amino acid racemases from *L. mesenteroides* with low temperature (30). *Leuconostoc* spp. capable of high D-amino acid production have also been isolated (24), therefore, the presence of a high abundance of *Leuconostoc* spp. may contribute significantly to the taste of *sake*, and need to isolate and culture *Leuconostoc* spp. found in this brewery for bacterial genome sequencing and detection of unique metabolism pathways.

### Nitrate-reducing bacteria are a major factor in microbial transitions

In this study, *Pseudomonas* spp. was detected in the middle stage, and the peak of nitrite reaction was observed on the 11th day (Fig. 1). On the other hand, previous studies on *Yamahai-moto* fermentation starters, which is a *sake* brewing recipe similar to *Kimoto*-style and lactic acid bacteria for maintaining the high acidity in the starter play an important role, reported cases where *Pseudomonas* spp. were not detected and no nitrite production was observed (5,31). It is considered that the genus *Pseudomonas* is easily lysed by exposure to alcohol due to its cell surface structure (32). Therefore, we believe that *Pseudomonas* spp. was detected until the middle stage, where the alcohol level rose relatively late, after day 20.

The structure of the microbiome changed significantly before and after the production of nitrite, suggesting that nitrate-reducing bacteria may also affect microbial transitions and other factors in the early stage (Fig. 1 and 2). This suggests that the presence or absence of nitrate-reducing bacteria such as *Pseudomonas* spp. may be one of the factors that cause differences in the microbial transitions in different breweries.

Taxonomic compositions of day 7 were markedly different between duplicates. Since the fermentation starter is fermented in a large tank and mixed with a stirring stick, it is considered that the microbial composition in the tank was not homogenous. In addition, 16S rRNA sequencing analysis possibly show subtle differences in each experiment due to experimental methods (33).

### The beneficial and harmful bacteria for *sake* brewing

*Bacillus* spp. have also been detected in *koji* and are known to be associated with the production of 4-vinyl guaiacol according to previous studies (9,34,35). In contract, the amylolytic activity of *Bacillus* spp. may play an important role in saccharification in the fermentation starter (9,36). The previous study that have examined the microbiome of *Kaburazushi* reported that *Staphylococcus* spp. and *Bacillus* spp. produce the components necessary for lactic acid bacteria growth (36). *Serratia* spp. have been identified by previous studies from *Yamahai-moto* (6), *Rahnella* spp. from *Yamahai* (25), and Chinese sauerkraut (37). *Hafnia-Obesumbacterium* spp. has been detected in brewer’s yeast along with *Rahnella* spp., suggesting that it may produce high pH beer by inhibiting fermentation reaction (38,39), it may also affect flavor in *sake* brewing. It is known that *Carnobacterium* spp. inhibits the growth of spoilage-associated bacteria and foodborne pathogens and has been estimated as useful bacteria for food preservations in various foods, including cold-smoked salmon, Ricotta Fresca cheese, and cooked ham (40–42). Therefore *Carnobacterium* spp. may contribute to preventing the growth of adventitious bacteria in *sake* brewing. A previous study (43) reported that *Methylobacterium* spp. were cultured in botrytized wine after 51 days of fermentation, suggesting that they may have also survived in the fermentation starter.

We found significant changes in relative abundances of these bacteria such as *Rahnella1, Serratia, Hafnia-Obesumbacterium* during the early to the late stages (Fig. 2). But These bacteria were not found during the early stage. Therefore, these bacteria may have been contaminated outside tanks during the middle stage (days 9-13). And these bacteria maintained high relative abundance and may have been grown in the fermentation starter for some time.

### Environmental microbiome mapping in the *sake* brewery

This study allowed us to detect the bacterial communities with phylogenetic composition during culturing of the fermentation starter comprehensively. In the early stage, a complex and varied microbiota is constructed as adventitious bacteria are contaminated by the built environments, tools, and raw materials used in the brewery. Since the *sake* brewing process is conducted in an open system, a variety of adventitious bacteria may contaminate the brewing’s *sake* from building environments, tools, and raw materials used in the brewery. Investigating the microbiome from architectural surfaces and tools in *sake* breweries may reveal the origin of adventitious bacteria in early *sake*. It has been suggested that these adventitious bacteria may affect the quality and taste of *sake*, so they need to be clarified (25,44). There are still many unexplained aspects of *Kuratuski* microorganisms, and we believe that this scientific elucidation of the traditional Japanese liquor will provide significant insights into food microbiology.

## Acknowledgment

Samples were collected by Mr. Kota Watanabe, Mr. Keitaro Nozaki, Ms. Chinami Fujita, Ms. Aimi Kurihara, Mr. Tsutomu Watanabe, Mr. Hiroaki Igarashi, Mr. Yuki Taguchi, and Ms. Mariko Kanazawa of the *Tsuchida Sake* Brewery. Amplicon sequencing was performed by GenomeRead Inc. We thank Morgenrot Inc. for providing the computational environment for the analysis. R.N. is a graduate student of Medical Innovation Program at Kyoto University and supported by JST SPRING, Grant Number JPMJSP2110. This study was funded by Tsuchida Sake Brewing Company.

## Data availability

The BioSample, DRA/SRA, and BioProject accession numbers for the sequence reported here are SAMD00513369-SAMD00513388, DRR393497-DRR393516, and PRJDB13924 respectively.

## Tables

TABLE S1. Total sequence reads per sample.

TABLE S2. Rarefaction curve of the Shannon index.

TABLE S3. Relative abundances of taxonomy.

TABLE S4. Statistical analysis of the microbial diversity of each brewery.

TABLE S5. Statistical analysis of the differences of each brewery.

